# Expression-based species deconvolution and realignment removes misalignment error in multispecies single-cell data

**DOI:** 10.1101/2021.04.04.438147

**Authors:** Jaeyong Choi, Woochan Lee, Jung-Ki Yoon, Sun Mi Choi, Chang-Hoon Lee, Hyeong-Gon Moon, Sukki Cho, Jin-Haeng Chung, Han-Kwang Yang, Jong-Il Kim

## Abstract

Although single-cell RNA sequencing of xenograft samples has been widely used, no comprehensive bioinformatics pipeline is available for human and mouse mixed single-cell analyses. Considering the numerous homologous genes across the human and mouse genomes, misalignment errors should be evaluated, and a new algorithm is required. We assessed the extents and effects of misalignment errors when using human and mouse combined reference data and developed a new bioinformatics pipeline with expression-based species deconvolution to minimize errors. We also evaluated false-positive signals for a species presumed to originate from ambient RNA of the other species and applied a computational method to remove them. Misaligned reads account for an average of 0.5% of total reads, but such reads were concentrated to few genes that were greatly affected. Human and mouse mixed single-cell data, analyzed using our pipeline, clustered well with unmixed data and showed higher k-nearest-neighbor batch effect test and Local Inverse Simpson’s Index scores than those derived from Cell Ranger. We also applied our pipeline to multispecies multisample single-cell library containing breast cancer xenograft tissue and successfully identified all samples using genomic array and expression. Moreover, diverse cell types in the tumor microenvironment were well captured. We present our bioinformatics pipeline for mixed human and mouse single-cell data, which can also be applied to pooled libraries to obtain cost-effective single-cell data. We also address major consideration points when analyzing multispecies single-cell data for other applications.

## Introduction

Single-cell RNA sequencing is a powerful method used to generate transcriptome with greater resolution [1], identify rare cells [2], and compare cell heterogeneity [3]. Single-cell technology also provides additional functionality such as the ability to discriminate the expression of human cells from mouse cells in human-mouse xenograft specimens or human-mouse cell coculture data [4, 5]. A simple method to process human and mouse mixed single-cell data is to align the data to human and mouse combined reference data (henceforth called “combined reference”). Cell Ranger (10× Genomics, Pleasanton, CA, USA) provides a combined reference and flags cellular barcodes where both human and mouse cells are presumably contained in a same droplet (henceforth called “cross-species doublet”) using read count distribution. However, similar sequences between each reference may cause human-originated reads to align to the mouse reference [6, 7]. In addition, because many mouse and human genes are homologous, some reads may have the best matching sequence in both references, leading to multi-mapped reads.

We measured the extent of misalignment error in single-cell data when using the combined reference to align human and mouse mixed single-cell data and developed a new realignment pipeline called Realignment-and Expression-based Multispecies deconvolution for Single cell (REMS) to minimize the errors. We also expanded and applied REMS pipeline to multisample, multispecies pooled single-cell library, enabling us to generate cost-effective single-cell libraries with minimal sequencing batch effects.

## Results

### Misalignment in the combined reference

To summarize the extent of misalignment in single-cell data, we compared the alignment of the combined reference and the human reference for several human single-cell datasets (**Table 1**). When aligning human-derived single-cell data to the combined reference, one would expect reads to align to the human part of the combined reference. Most reads indeed aligned to human genes, but depending on the data type and sequencing throughput, 3,000–480,000 reads misaligned to the mouse genes. More importantly, when comparing data analyzed using the combined reference and the human reference, 13,000–300,000 fewer reads aligned to human genes in the combined reference than in the human reference. Misalignment errors were more prominent in tumor samples, and errors were also found on aligning mouse-derived single-cell data to the combined reference. Although all incorrectly aligned reads accounted for only 0.3%–1% of total reads, these reads were concentrated to few genes, leading to strong false signals (**Fig 1**). For example, 30%–65% of reads of *HNRNPA3* were lost in human cells, and nearly all reads (99.8%) of *Erf1* were lost in mouse cells. Genes related to RNA binding and human leukocyte antigen genes had high rates of misalignment errors. Interestingly, the genes affected by misalignment and the degree of misalignment were not identical between samples and tissues. Genes with read gain and genes with read loss did not overlap for human and mouse data, suggesting that misalignment was not a simple switch-of-position.

**Table 1.** Misaligned reads in the combined reference.

| Data name | Species | Tissue | Cell count | Matching reference | Combined reference |  | misaligned reads % |
| --- | --- | --- | --- | --- | --- | --- | --- |
|  |  |  |  | Total reads | Reads aligned to correct reference | Reads misaligned to other reference |  |
| placentaD121 | human | placenta | 2,522 | 35,877,521 | 35,828,222 | 11,252 | 0.17 |
| liverP1 | human | liver | 957 | 3,179,327 | 3,165,784 | 3,243 | 0.53 |
| deciduaD12 | human | decidua | 1,963 | 25,147,111 | 25,122,973 | 6,205 | 0.12 |
| 10x_pbmc | human | peripheral blood mononuclear cells | 9,295 | 93,171,679 | 93,027,903 | 175,980 | 0.34 |
| kidneyF21 | human | kidney | 3,399 | 21,326,290 | 21,266,741 | 77,660 | 0.64 |
| 10x_hodgkin | human | hodgkin lymphoma | 2,502 | 19,341,273 | 19,305,209 | 22,330 | 0.30 |
| 10x_SCC | human | small cell cancer | 4,739 | 72,819,855 | 72,583,115 | 482,564 | 0.99 |
| 10x_lungcancer | human | lung cancer | 4,481 | 50,316,076 | 50,165,228 | 253,394 | 0.80 |
| 10x_adenocarcinoma | human | adenocarcinoma | 3,599 | 44,820,075 | 44,678,244 | 268,975 | 0.92 |
| 10x_NSCLC | human | non small cell lung cancer | 6,640 | 69,706,720 | 69,466,298 | 478,684 | 1.03 |
| 10x_mbrain | mouse | brain | 8,096 | 103,833,090 | 103,524,395 | 38,599 | 0.34 |
| mcolonHC1 | mouse | colon | 1,204 | 8,810,265 | 8,782,774 | 2,440 | 0.34 |

**Fig 1.**
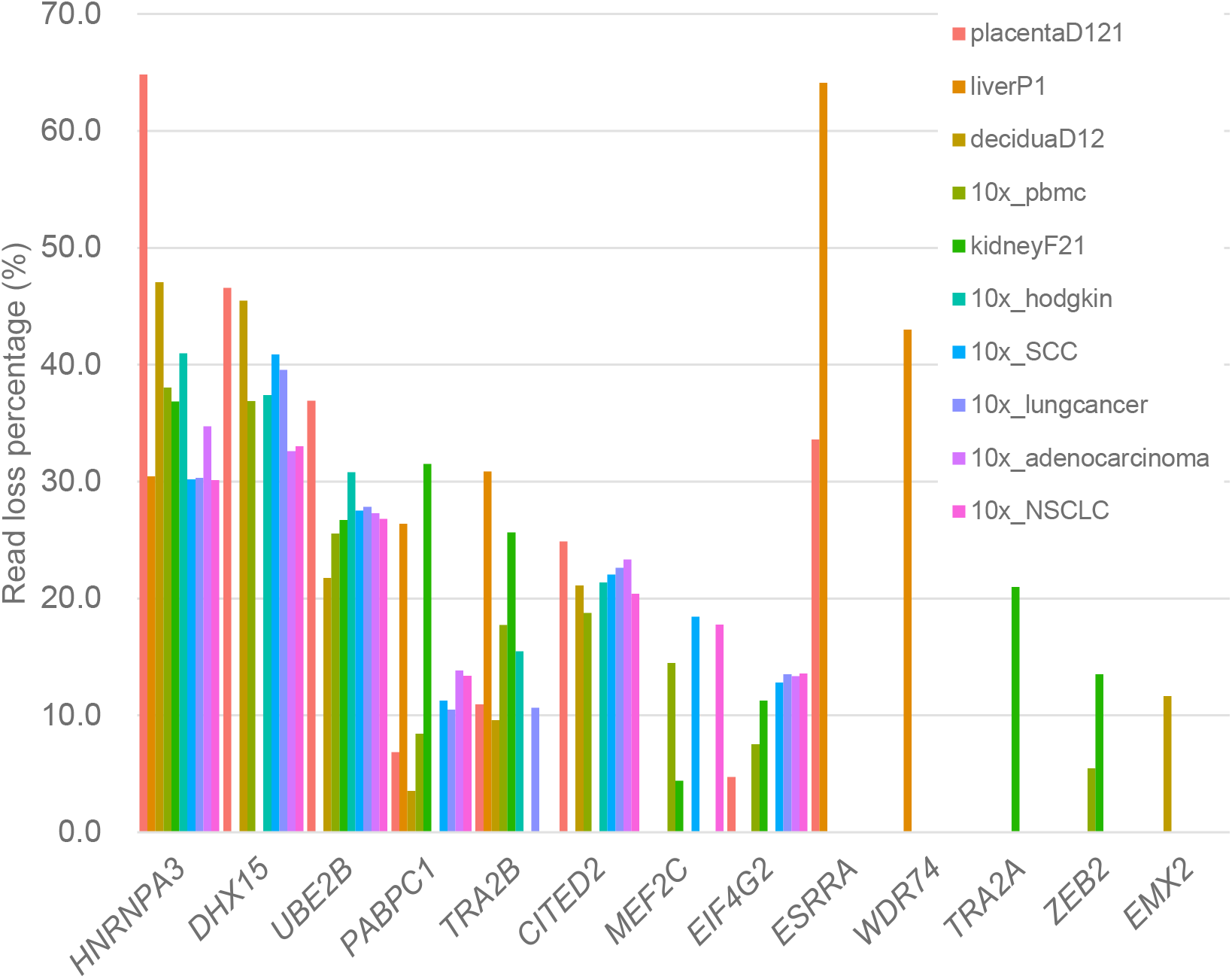
Read loss percentage of the most misaligned genes.

### Mixed data analysis pipeline

To assess the misalignment error in human and mouse mixed single-cell data, we generated both mixed and separate libraries of human lung tissue, human gastric organoids, and mouse intestinal organoids. Our mixed data also had misalignment error when we aligned the data to the combined reference (**S1 Table**). The error was higher, possibly owing to ambient RNA contamination and undetected doublets. To overcome this issue, we generated REMS pipeline to remove doublets and ambient RNA signals for error-free downstream analysis (**Fig 2A**).

**Fig 2.**
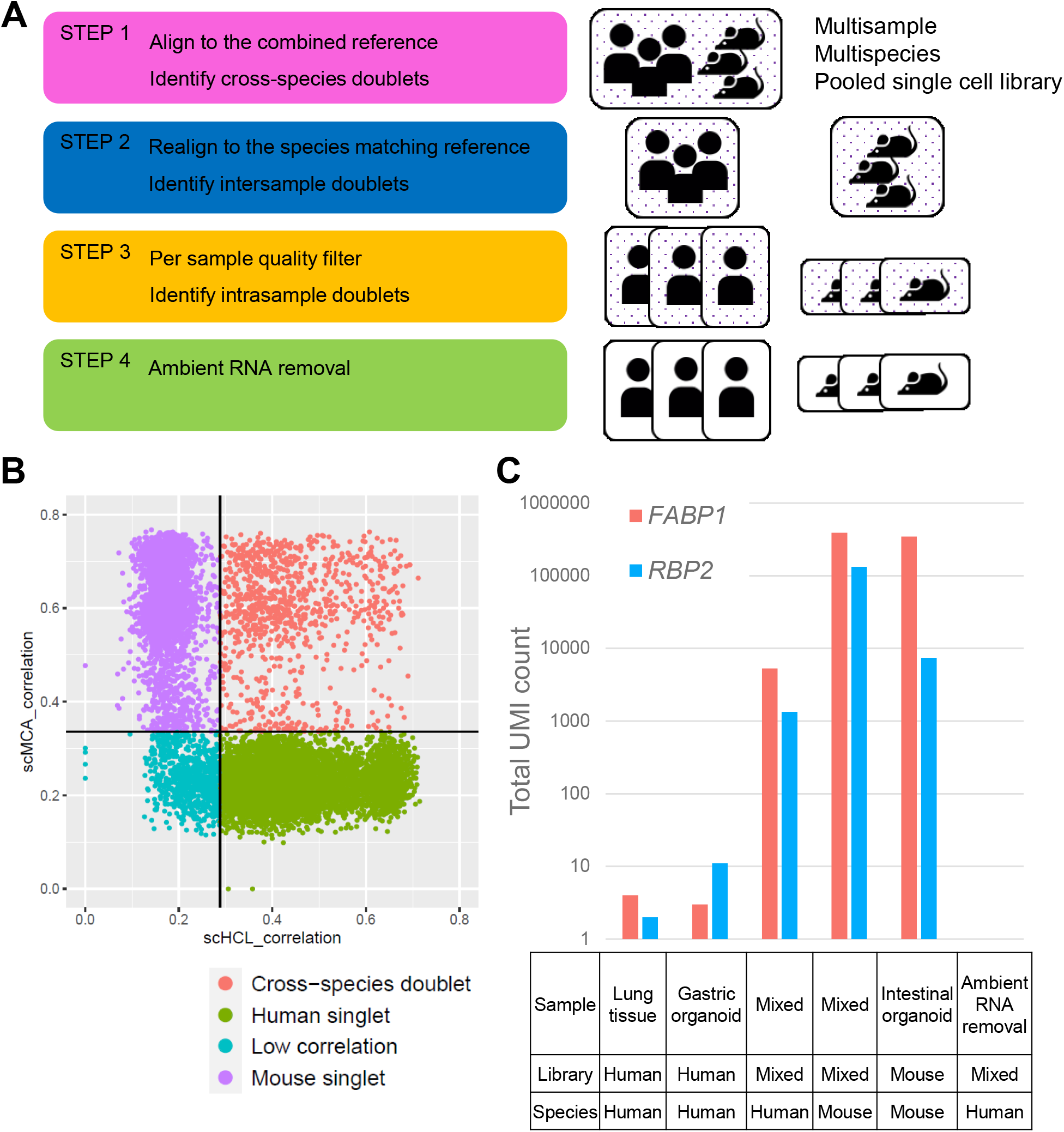
(A) REMS pipeline schema. (B) Identification of cross-species doublets by expression correlation. Black horizontal and vertical lines are thresholds for doublet identification. (C) Misaligned reads generated from ambient RNA of different species.

First, we aligned mixed data to the combined reference and identified species and cross-species doublets. Then, we applied a minimum quality filter during the first step, because the quality distribution for human and mouse cells may differ widely. To identify cross-species doublets, we used transcriptome-wide expression levels to calculate the correlation to known single-cell data (**Fig 2B**). As misalignment occurs for a proportion of genes, the overall expression pattern would be more robust to error. True cross-species doublets would show expression patterns of both human and mouse cells. We note that although barcodes with low read counts did not reach the threshold for doublet identification, >90% of all cells with a low correlation value were filtered in future steps.

After removing cross-species doublets, we realigned the data to each species matching reference and selected barcodes of the corresponding species. When multiple identities were pooled, we used a variant-based deconvolution method to assign identity to each droplet and detect intersample doublets generated from different cell identities. After doublet filtration, we applied quality filters for each identity to account for sample level quality variation, and then used a doublet detection algorithm based on artificial doublet generation to identify intrasample doublets originating from the same identity.

Although we used species matching reference to remove misalignment errors, human singlet cells sequenced from mixed data contained reads that were not expressed in separately sequenced human cell data, namely *FABP1* and *RBP2* (**Fig 2C**). Reads aligning to these genes were found in ambient RNA of the mixed data and in separately sequenced mouse data aligned to the human reference, but not in ambient RNA of separately sequenced human data. These findings suggested that mouse RNA was introduced into human cell containing droplets in the form of ambient RNA, and it misaligned to human genes.

Since ambient RNA generated reads from the other species misalign similarly in case of empty droplets and human cells, methods that remove ambient RNA signals based on expression levels of empty droplets will remove ambient RNA signals generated from both species. We strictly removed ambient RNA using a contamination rate derived by adding contamination level calculated from each species with an additional 0.05. After ambient RNA removal, all false signals in *FABP1* and *RBP2* were removed.

To examine the downstream effect of misalignment, we compared the integration level between mixed and unmixed data for the data generated by Cell Ranger and that generated by REMS pipeline (**S1 Fig**). A total of 98.9% of cells clustered to identical cell types for each dataset, which reflects the small effect of misalignment error in overall expression. However, drastic changes was found in integration metrics. Even after batch effect correction using Harmony, a large difference in the k-nearest-neighbor batch effect test (kBET) was found for gastric organoid cells, overall lung tissue, macrophages, ciliated cells, and neutrophils (**Table 2**). REMS pipeline also had higher Local Inverse Simpson’s Index (LISI) values for most samples and celltypes.

**Table 2.** Comparison of integration metrics between Cell Ranger and REMS pipeline. 0 indicates no integration and 1 indicates full integration. Integration metrics for each sample and celltypes with ≥100 cells were calculated.

| Data type | kBET |  |  | LISI |  |  |
| --- | --- | --- | --- | --- | --- | --- |
|  | CellRanger | REMS | Difference | CellRanger | REMS | Difference |
| mIO04 | 0.802 | 0.800 | -0.001 | 0.916 | 0.927 | 0.011 |
| hGO03 | 0.051 | 0.859 | 0.807 | 0.907 | 0.909 | 0.003 |
| hLT02 | 0.246 | 0.387 | 0.141 | 0.868 | 0.876 | 0.008 |
| Macrophage | 0.620 | 0.952 | 0.333 | 0.814 | 0.905 | 0.091 |
| Ciliated | 0.321 | 0.506 | 0.184 | 0.842 | 0.868 | 0.026 |
| Neutrophil | 0.800 | 1.000 | 0.200 | 0.929 | 0.937 | 0.007 |
| Monocyte | 0.904 | 0.985 | 0.081 | 0.921 | 0.928 | 0.006 |
| Endothelial_1 | 0.981 | 1.000 | 0.019 | 0.884 | 0.908 | 0.024 |
| Mast | 0.963 | 0.973 | 0.010 | 0.942 | 0.939 | -0.004 |
| Endothelial_2 | 1.000 | 1.000 | 0.000 | 0.936 | 0.923 | -0.013 |
| Alveolar Type 2 | 0.969 | 0.962 | -0.007 | 0.853 | 0.946 | 0.093 |

We then applied our pipeline to another library containing breast cancer patient-derived xenograft tissue and multiple organoids. Because the gene expression of tumor cells does not resemble that of normal cells, we additionally used copy number clustering in the cross-species doublet identification step. From 23,642 initial cells called by Cell Ranger, we identified 6,738 human cells with six different identities and 7,580 mouse cells with two different identities (**Fig 3A and 3B**). Human tumor cells (xBC42) had a high level of aneuploidy, and the copy number pattern was identical to that in exome sequencing-derived data (**S1 Fig**). Four of the human cell identities were lung organoids: three normal organoids (hLO13, hLO18, and hLO19) clustered evenly while one treated with interleukin 13 (hLO11) had a distinct cluster. The remaining human cell identity was breast organoid (hBO25); fewer cells than expected had this identity. All human identities had matching genotypes to genomic array data. Two mouse identities are microenvironment of xenograft sample (xME42) and mouse pancreatic islet organoid (mPO31). The diverse tumor microenvironment was well captured. Moreover, two groups of macrophages, endothelial, stromal, and muscle cells as well as normal alveolar, luminal, and myoepithelial cells were identified.

**Fig 3.**
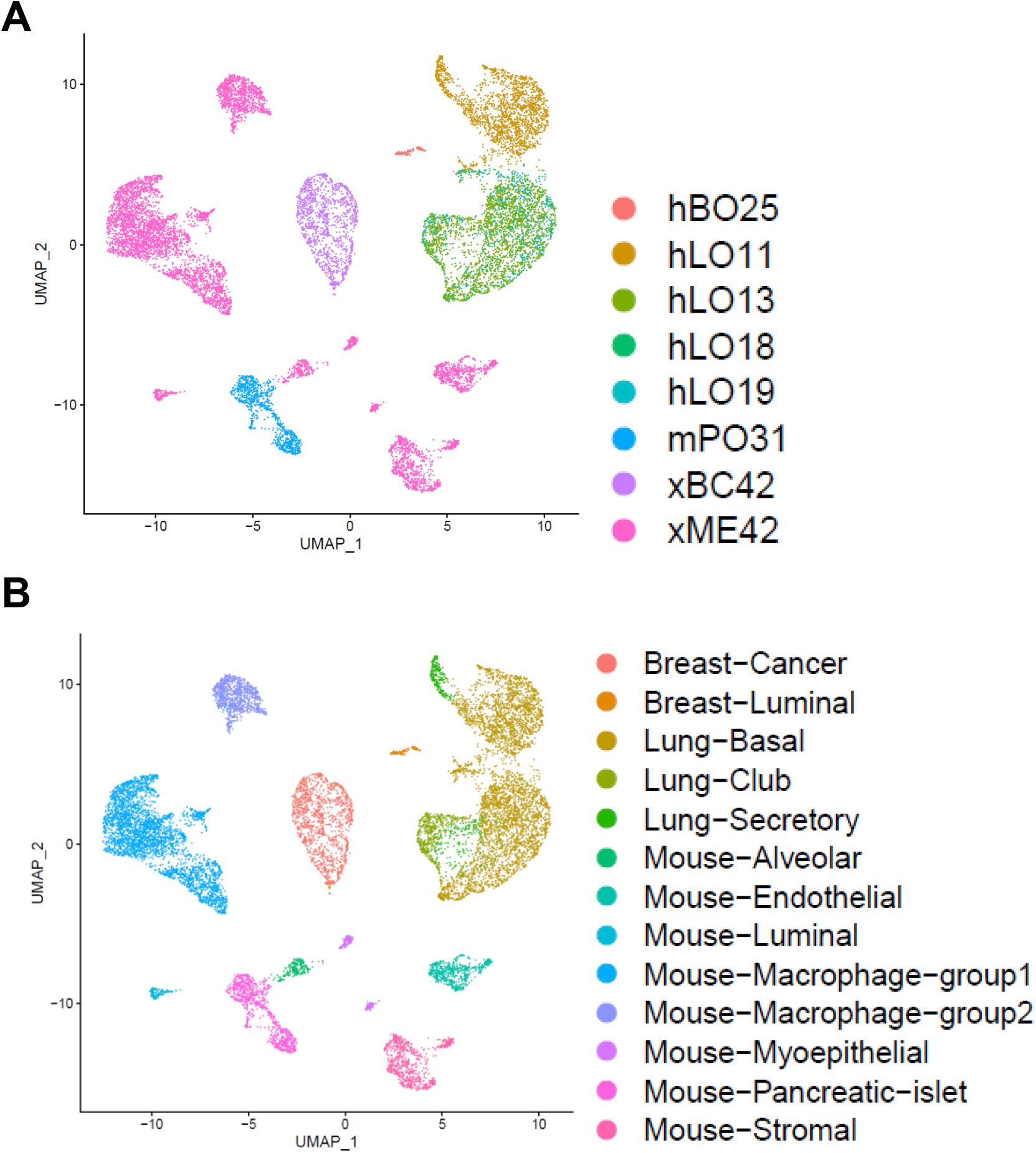
Cell identity (A) and cell type (B) after REMS pipeline.

## Discussion

When analyzing multispecies single-cell data, misalignment errors greatly affect the results for some genes. The errors were not identical across tissues or cell types, and some cells had >10% of reads misaligned to genes of other species. Due to misalignment, cross-species doublet identification via read count or read majority [5, 8] may lead to inaccurate calls. In addition, ambient RNA from other species can generate strong false-positive signals, which may disturb downstream analysis.

To cope with misalignment error, we highlight three major consideration points. First, the origin of reads in a single droplet should be identical. Identifying species for each droplet while confidently removing cross-species doublet is required. Second, realigning each droplet to the corresponding reference is required to minimize misalignment error and multiple-mapping events. Third, ambient RNA should be adequately estimated and should be removed.

We developed a novel pipeline called REMS using expression-based species deconvolution. Our method can handle pooled library and can be used to generate cost efficient single-cell libraries. Overloading a single-cell library to target 40,000 droplets, accompanied by adequate doublet removal can reduce the cost of library generation to as much as 1/10 per singlet. Our pipeline is not restricted to any specific tool, and one can use other methods as long as they can minimize misalignment error and adequately remove ambient RNA signals.

Multiple deconvolution methods with or without external treatment are available for multisample pooling. CITE-seq [9] uses oligo-tagged antibodies that target commonly expressed cell surface proteins. MULTI-seq [10] uses a similar method but with lipid-tagged indices that merge into the membrane of cells or nuclei. Variant-based method uses natural genetic variation sequenced in RNA reads to identify each cell [4, 11].

Most methods cannot be applied to xenograft samples or human and mouse cocultured samples, wherein mouse and human cells cannot be separated. Lipid-tagged barcodes are agnostic to species and currently there is no variant-based deconvolution tool that handles multispecies data. Cell barcode antibodies that target human cells or mouse cells separately or a conventional cell sorting technique can be used to separate mixed cells [12]. However, both methods need additional processing, which induces stress in the cells. We attempted to establish an unbiased approach using minimal treatment.

During the development of REMS, we found that some droplets had high mitochondrial gene percentage of the other species, but they were not detected as cross-species doublets using multiple detection methods. The mitochondrial gene percentage in these droplets was much higher than that in ambient RNA, and the possibility of ambient RNA owing to its expression was low. We hypothesize that small cell debris may get mixed with droplets and generate partial cross-species doublets that may be much harder to detect than cross-species doublets. Cell debris from the same species may also form partial doublets and may be mistaken as novel cell types or transient cell states. In this study, cells were handled swiftly by experienced personnel to minimize stress and all cells with high mitochondrial percentage were stringently filtered.

In summary, we identified misalignment error while using the combined reference and present our realignment pipeline, which is robust for misalignment to generate error-corrected gene matrix for downstream analysis. Tools for multispecies single-cell data are lacking, and we encourage researchers to optimize our pipeline strategy using advanced computational methods.

## Materials and methods

### Tissue preparation

Normal human tissues were obtained from early-stage lung, breast, and stomach cancer surgery. Pathologists confirmed the absence of cancer cells. Three samples of human lung tissues were obtained from tissue donations of three patients with idiopathic pulmonary fibrosis who underwent lung transplantation. A bronchoscopic brushing sample was collected from healthy bronchus obtained from a patient who received bronchoscopy. Furthermore, a breast cancer patient-derived xenograft mouse was sacrificed 6 weeks after tumor injection. All studies were conducted in accordance with the Declaration of Helsinki and the Guidelines for Good Clinical Practice and performed after receiving informed consent from each patient. Mouse normal islet and intestine tissues were obtained from a 12-week-old C57BL/6 mouse. The institutional review board of the Seoul National University Hospital granted permission to perform this study (2008-065-1148, 1602-108-742, 1909-110-1066, 1805-151-948, and 1402-054-555).

Fresh human tissues were treated using the Multi Tissue Dissociation Kit 1 (Miltenyi Biotec, Bergisch Gladbach, Germany) along with the gentleMACS Octo Dissociator (Miltenyi Biotec) and 37C_Multi_A_01 program. The brushing sample was centrifuged at 400 g for 5 min at 4°C and the pellet was resuspended in Advanced DMEM/F12 (Gibco, Dublin, Ireland) with Glutamax (Gibco), HEPES (Gibco), Penicillin-Streptomycin (Gibco) (ADF^+++^), and 1 mg/mL collagenase IV (Sigma, St. Louis, MO, USA), followed by digestion for 15 min at 37°C. Resected xenograft tumor tissue was dissociated into a single cell suspension using the Tumor Dissociation Kit 1, human (Miltenyi Biotec) along with the 37C_h_TDK_3 program. Mouse islet cells were purified using Ficoll-Paque (Cytiva, Marlborough, MA, USA) and digested using TrypLE™ (Gibco) for 15 min at 37°C. Mouse intestinal crypt cells were isolated using the Gentle Cell Dissociation Reagent (STEMCELL Technologies, Vancouver, Canada) for 20 min at room temperature followed by squeezing using a microscope cover glass.

Each digested tissue (without intestinal crypt cells) suspension was strained using a 70-μm filter. The suspension was then centrifuged at 400 g for 5 min at 4°C with 10 mL of ADF^+++^. The pellet was resuspended in 1 mL of 1× Red Blood Cell Lysis Solution (Miltenyi Biotec) for 10 min at room temperature for lysis of erythrocytes, followed by the addition of ADF^+++^ and centrifugation at 400 g for 5 min. The pellet was then resuspended in an appropriate volume of ADF^+++^ and kept on ice for organoid culture or library generation.

### Organoid culture

After digestion and erythrocyte lysis, the cell pellets were seeded in Corning® Matrigel® Growth Factor Reduced Basement Membrane Matrix (Corning, Corning, NY, USA), solidified for 15 min at 37°C, and cultured on each organ-specific media for human lung [13], human breast [14], human gastric [15], mouse islet [16], and mouse intestine [17] cells. The media was changed every 2–3 days and the organoids were subcultured every 2–3 weeks depending on the density and growth rates of organoids.

### Single-cell RNA sequencing

We performed single-cell RNA sequencing according to the standard 10× Genomics 3′ V3.1 chemistry protocol (10× Genomics). Before loading cells on the G chip, cells were counted to determine the cell concentration. The number of loading cells per library was calculated via the multiplexing cost calculator (satijalab.org/costpercell/). The libraries were sequenced using the NovaSeq 6000 (Illumina, San Diego, CA, USA) platform S4 with paired-end 100 bp.

### Data download

Human decidua (D12) [18], placenta (D121), liver (P1) [19], kidney (F41) [20], and mouse colon (HC1) [21] single-cell data were downloaded from the Human Cell Atlas Data Portal (data.humancellatlas.org). Human blood, Hodgkin lymphoma, pooled cancer, and mouse brain data were downloaded from 10× Genomics single-cell gene expression datasets (support.10xgenomics.com/single-cell-gene-expression).

### Assessment of misalignment

We aligned the data to species matching reference data (refdata-gex-GRCh38-2020-A or refdata-gex-mm10-2020-A) using Cell Ranger 4.0.0 (10× Genomics). Seurat v3.2.3 [22] was used to filter barcodes by low total read count, low expressed gene count, and high mitochondrial gene percentage. We extracted these cellular barcodes from the data aligned to the combined reference (refdata-gex-GRCh38-and-mm10-2020-A) and compared the alignment result. We defined misaligned read as the following: 1) read aligned to a gene of the other species or 2) read difference between the combined reference and the species matching reference. We defined “misaligned gene” as the following: 1) average read count higher than 0.1 per cell and 2) >10% difference in the read count between the species matching reference and the combined reference. Pathway analysis for misaligned genes was performed using the web server g:Profiler [23].

### REMS pipeline step 1: Alignment and prefilter

We aligned human and mouse mixed single-cell data to the combined reference using Cell Ranger 4.0.0. Seurat v3.2.3 [22] was used to filter cells, normalize expression, cluster, and visualize data. The EmptyDrops methods [24] was used to identify droplets that had an expression pattern similar to that of ambient RNA. We selected barcodes with expressed gene count ≥200 in the human reference and/or mouse reference. We removed barcodes if their percentage of mitochondrial genes was twice or higher than the percentage of mitochondrial genes in ambient RNA; this was performed separately for human and mouse genes.

### REMS pipeline step 2: Expression-based cross-species doublet identification

For data containing nontumor cells, we used the single cell Human Cell Landscape (scHCL) [25] and single cell Mouse Cell Atlas (scMCA) [26] correlation values to estimate cross-species doublets. The cutoff was set to 99% confidence interval of median normalized correlation value for human and mouse cells. Barcodes with both high correlations to human cells and mouse cells were classified as cross-species doublets. Other barcodes were classified as human cells or mouse cells according to read majority and correlation value. We removed barcodes with discrepancies. When expression for each species could be clearly separated by clustering, minor species in each cluster were removed.

For data containing tumor cells, we used copy number cluster result from copyKAT [27] in addition to correlation values to identify tumor cells with low correlation value. We defined aneuploid barcodes as tumor cells. Barcodes with both valid copy number result and mouse read majority were additionally classified as cross-species doublets.

### REMS pipeline step 3: Species matching reference realignment

After species identification, we realigned data to the human reference or mouse reference, and selected species matching barcodes. When multiple samples were pooled, we used Souporcell [11] to identify intersample doublets. We selected singlets identified using Souporcell, and when the expression for each sample could be clearly separated, minor identities in each cluster were removed. Normal cells with noisy copy number segments were also removed.

For each sample, we applied a quality filter separately to account for sample variation. Cells with low total read count, low expressed gene count, and high mitochondrial gene percentage were filtered. Separately filtered barcodes were merged and DoubletFinder [28] was used to identify intrasample doublets.

### REMS pipeline step 4: Ambient RNA removal

SoupX [29] was used to estimate the ambient RNA expression from barcodes with ≤100 reads. We calculated the contamination rate in human and mouse cells separately. We additionally added 0.05 to the contamination rate to stringently remove ambient RNA signals. Additional information regarding this bioinformatics pipeline can be found at snumrc.snu.ac.kr/gmi/en/community/gallery?md=v&bbsidx=121.

### Result comparison

Harmony [30] was used to remove batch effect from mixed and separately sequenced libraries. Cell type was identified by the consensus of scHCL correlation for each cluster. We used the kBET [31] and LISI [30] with top 20 components computed with Harmony. We calculated integration metrics for each sample and celltypes with ≥100 cells. Metrics for T cell, Granulocyte, and Club_Basal cluster were not calculated due to heterogeneous subtypes within each cluster. Each metric was normalized to a 0–1 scale, 0 indicating no integration and 1 indicating full integration.

### Genomic array

DNA was extracted from tissue or cultured organoids using DNeasy Blood & Tissue Kit (Qiagen, Germantown, MD, USA). Axiom Korea Biobank Array (ThermoFisher Scientific, Waltham, MA, USA) [32] was used to genotype samples. PLINK v1.0.9 identity-by-descent [33] was used to confirm single-cell deconvoluted sample identity.

### Data availability

Mixed and separately sequenced single-cell data are available at Sequence Read Archive under BioProject number PRJNA719675. Xenograft and lung organoid single-cell data are not publicly available due to institutional review board restrictions and are available from the corresponding author on reasonable request.

## Funding

This work was supported by a National Research Foundation of Korea grant funded by the Korean government (2020R1A2C3012524) and by a Basic Science Research Program through the National Research Foundation of Korea funded by the Ministry of Education (2020R1A6A1A03047972).

## Competing interests

The authors have declared that no competing interests exist.

## Figure Legends

**S1 Fig. UMAP of human lung cells processed using Cell Ranger annotated by library (A) and cell type (B). UMAP of human lung cells processed using REMS pipeline annotated by library (C) and cell type (D)**.

**S2 Fig. Copy number from single-cell and exome data**.

## Notes

https://snumrc.snu.ac.kr/gmi/en/community/gallery?md=v&bbsidx=121

### Competing Interest Statement

The authors have declared no competing interest.

