## Supplementary figures and images for "Expression-based species deconvolution and realignment removes misalignment error in multispecies single-cell data"

### Supplemental Figure 1

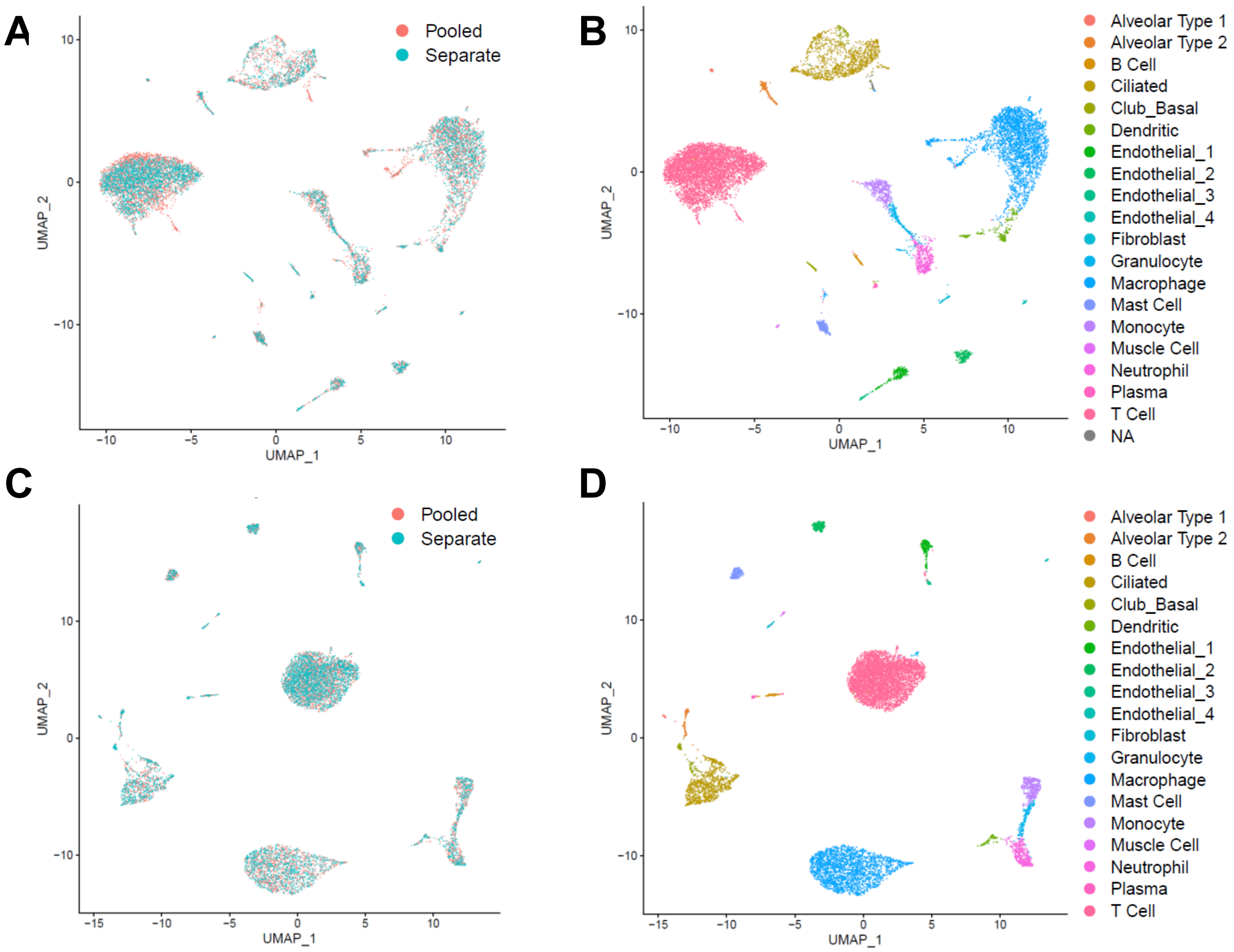

### Supplemental Figure 2

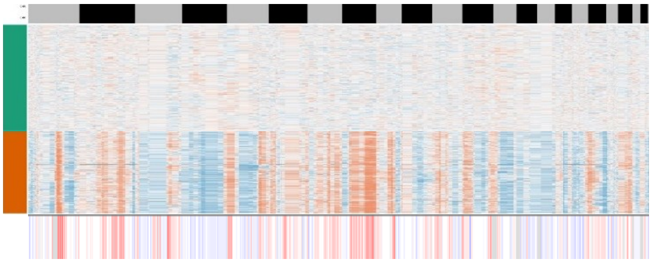
